# The Prolonged Terminal Phase of Human Life Induces Survival Response in the Skin Transcriptome

**DOI:** 10.1101/2023.05.15.540715

**Authors:** Ahmed S Abouhashem, Kanhaiya Singh, Rajneesh Srivastava, Sheng Liu, Shomita S Mathew-Steiner, Xiaoping Gu, Sedat Kacar, Amit Hagar, George E. Sandusky, Sashwati Roy, Jun Wan, Chandan K Sen

**Author notes:** Corresponding Authors: Chandan K. Sen, PhD;, IB454, 975 W Walnut Street, Indianapolis, IN-46202, USA, Kanhaiya Singh, PhD;, IB444, 975 W Walnut Street, Indianapolis, IN-46202, USA. Equal contribution (Ahmed S Abouhashem, Kanhaiya Singh).

## Abstract

Human death marks the end of organismal life under conditions such that the components of the human body continue to be alive. Such postmortem cellular survival depends on the nature (Hardy scale of slow-fast death) of human death. Slow and expected death typically results from terminal illnesses and includes a prolonged terminal phase of life. As such organismal death process unfolds, do cells of the human body adapt for postmortem cellular survival? Organs with low energy cost-of-living, such as the skin, are better suited for postmortem cellular survival. In this work, the effect of different durations of terminal phase of human life on postmortem changes in cellular gene expression was investigated using RNA sequencing data of 701 human skin samples from the Genotype-Tissue Expression (GTEx) database. Longer terminal phase (slow-death) was associated with a more robust induction of survival pathways (PI3K-Akt signaling) in postmortem skin. Such cellular survival response was associated with the upregulation of embryonic developmental transcription factors such as *FOXO1*, *FOXO3*, *ATF4* and *CEBPD*. Upregulation of PI3K-Akt signaling was independent of sex or duration of death-related tissue ischemia. Analysis of single nucleus RNA-seq of post-mortem skin tissue specifically identified the dermal fibroblast compartment to be most resilient as marked by adaptive induction of PI3K-Akt signaling. In addition, slow death also induced angiogenic pathways in the dermal endothelial cell compartment of postmortem human skin. In contrast, specific pathways supporting functional properties of the skin as an organ were downregulated following slow death. Such pathways included melanogenesis and those representing the skin extracellular matrix (collagen expression and metabolism). Efforts to understand the significance of death as a biological variable (DABV) in influencing the transcriptomic composition of surviving component tissues has far-reaching implications including rigorous interpretation of experimental data collected from the dead and mechanisms involved in transplant-tissue obtained from dead donors.

## Introduction

Post-mortem tissue samples are a valuable resource for biological research. Specifically, use of post-mortem material is crucial for studying the patterns of gene expression underlying tissue specificity within individuals, as sampling such tissues from living individuals would be challenging. Samples obtained post-mortem are a valuable source of material for studies requiring organs and tissues difficult to obtain, or those where it is not feasible to study and manipulate them in living organisms. Hence, understanding the impact of death in tissues is essential for the rigorous interpretation of post-mortem gene expression levels as a proxy for *in vivo* living physiological conditions. Agonal factors refer to the manner of death and the terminal state before death. Death has been classified as slow, intermediate, fast from natural causes, and violent fast^1^. Terminal states include coma, inadequate oxygen, fever, infection, and artificial respiration^2, 3^. Previous works have shown that messenger RNA (mRNA) levels are sensitive to agonal factors ^4,5^and related changes in pH of the affected tissues^6^. Some studies include an agonal factor score to account for specific agonal conditions and terminal states per individual^7, 8^. However, the basic design typically retains the following shortcomings: (i) studies focused primarily on the correlation of agonal factors to aspects such as the RNA integrity number, tissue pH, and whole gene expression pattern, neglect to identify the mechanistic underpinnings underlying agonal related regulation; (ii) such studies^6–9^ often roughly combine various terminal states together, thereby overlooking the potential interactions or conflicts between them; and, (iii) the analytical approach is often limited including Pearson correlation and differential gene expression neglecting gene co-expression relationships.

Contrary to other bodily organs such as the brain and heart, the skin tissue maintains its integrity for several hours after death^10^. No obvious histological changes were identified in either epidermal or human dermal skin layer in the first 18h post-mortem^10^. Thus, employing skin as the organ of choice for this study, this work sought to test the hypothesis that human death is followed by the induction of specific molecular signaling processes which support survival of component skin cells. Mounting of such adaptive response is time-dependent and thus plays out effectively during natural slow death but not during fast death. Thus, the post-mortem skin transcriptome was investigated under conditions of slow and fast death. To address this objective, RNA sequencing data of 701 sun-exposed human skin samples from the Genotype-Tissue Expression (GTEx) database were mined^11^. This work identifies differentially expressed genes (DEGs), pathways and biological processes in sun-exposed skin samples from dead humans with prolonged terminal phase of life (slow death) compared to those with short terminal phase of life (fast death).

## Results

### Longer terminal phase (slow death) was associated with robust induction of survival pathways in the postmortem skin

mRNA sequencing data from sun-exposed skin tissue present in the GTEx project v8 were obtained^8^ (**Supplementary Table 1**). These skin samples were classified into five categories based on the 4-point Hardy scale^8^. This classification was utilized to identify the effect of the prolonged terminal phase of life on skin transcriptome as in the case of slow death-type. A total of 701 samples included 28 samples of fast violent death type (a score of 1; terminal phase: <10 mins), 192 samples of fast unexpected death type (a score of 2; terminal phase: <1h), 39 samples of intermediate death type (a score of 3; terminal phase: 1 – 24h), 81 samples of slow death type (a score of 4; terminal phase: >24h) and 352 samples of cases on a ventilator immediately before death (a score of 0). Nine samples had no scores assigned (**Fig. 1A-B; Supplementary Table 1**). Samples assigned to different death conditions showed distinct distribution in the reduced space using Uniform Manifold Approximation and Projection (UMAP) in which samples from slow death condition were scattered mainly in the middle, while samples from fast death were scattered mainly at the bottom of the plot (**Fig. 1B; Supplementary Fig. S1A**). Additionally, the percentage of variance explained by death type was higher than variance explained by age, sex, RNA quality or ischemic time (**Supplementary Fig. S1B)**.

**Figure 1.**
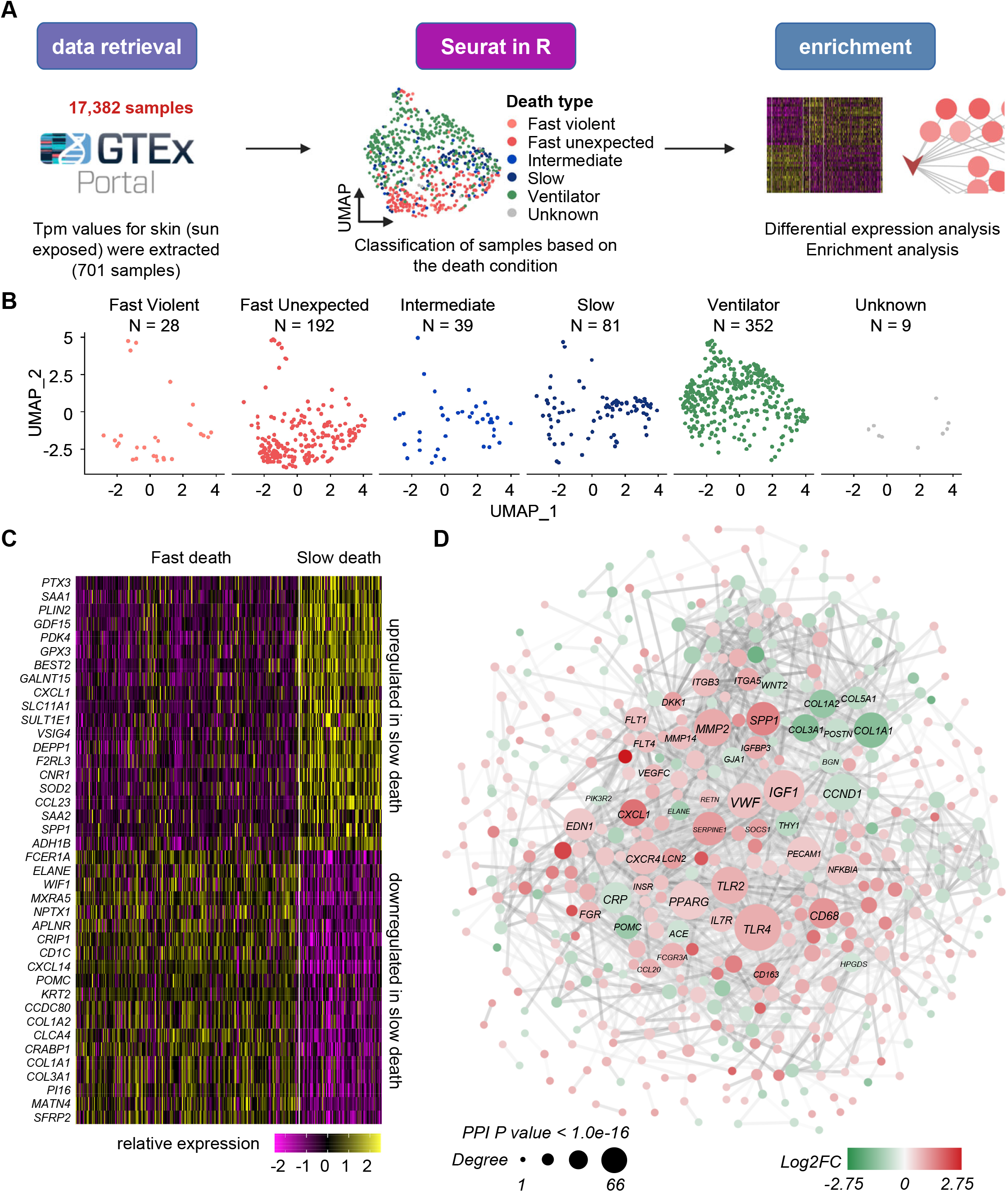
Analysis of transcriptomic data in human post-mortem sun-exposed skin based on type of death. (**A**) The analysis workflow. (**B**) UMAP plot showing total human skin samples included in this study (701 samples). Each sample is represented as a dot. (**C**) Heatmap representing the top 20 upregulated and top 20 downregulated differentially expressed protein coding genes (DEG) in skin cells of slow death-type (Hardy scale 4) cases compared to fast (Hardy scale 1 and 2) death-type cases. Gene values are log normalized; see methods. (**D**) Protein-protein interaction network between DEG with log2FC >±0.5 and adjusted p value < 0.01, Wilcoxon Rank Sum test. Node color represents log2FC. Node size represents node degree (number of protein-protein interactions). Nodes with degrees > 20 were labeled. The protein-protein interaction network enrichment has p value < 1.0e-16.

The present study catalogues transcriptomic differences between skin cells from fast and slow death types. Ventilator cases were excluded as the death type of the individual encountered was not documented. Comparing gene expression level from slow death-type cells (Hardy scale score = 4) *versus* fast death-type samples (a hardy scale score of 1 and 2) resulted in identification of 1,582 differentially expressed genes (DEG) (adjusted p value using Bonferroni method < 0.01, **Supplementary Table 2**). Among them, 344 genes were upregulated, and 183 genes downregulated (cutoff log2FC >± 0.5) (**Fig. 1C; Supplementary Table 2**). Among the identified DEG, a protein-protein interaction (PPI) network of 451 genes (nodes) was identified connected with 1,892 edges (p value <1.0e-16). The hub genes ranked by the highest degree in the PPI network included *TLR4*, *IGF1*, *PPARG*, *MMP2*, *TLR2*, *CCND1*, *COL1A1*, *VWF* and *PECAM1* (**Fig. 1D; Supplementary Table 2**). To investigate whether the observed transcriptomic changes in slow death were sex-specific, additional analysis was performed. Comparison between males and females with respect to death type identified a total of 100 DEGs (48 upregulated and 52 downregulated) in slow death samples irrespective of the sex (adjusted p value <0.05 and log2FC cutoff >±0.5, **Supplementary Table 3)**. Among the unique DEG in females, 6 were upregulated and 32 were downregulated in slow death skin as compared to fast death (adjusted p value < 0.05). The upregulated genes in female slow death skin included *SAA4*, *AP001033.2*, *KLHL24*, *YTHDC1*, *RSL1D1* and *C20orf141* (**Supplementary Fig. S1C-D**). Although these genes altogether did not result in any specific pathway enrichment, *YTHDC1,* was an interesting candidate found to be upregulated in skin cells from females with slow death type. *YTHDC1*, a nuclear mRNA reader, is responsible for RNA modification N6-methyladenosine (m6A) regulation and maintenance of embryonic stem cells^9^. That depletion of YTHDC1 leads to early embryonic lethality in mice, exhibit the critical role of this gene in context of development biology^12, 13^. The regenerative functions of YTHDC1 have been documented in the setting of acute muscle injury^14^. Post-injury, induction of YTHDC1 augmented skeletal muscle stem cell activation and proliferation. In contrast, inducible YTHDC1 depletion decimated stem cell regenerative capacity^14^. Furthermore, YTHDC1 is known to regulate mouse embryonic stem cell self-renewal by recognizing m6A- modified LINE1 RNA^15^.

Next, to understand the biochemical processes associated with the skin of slow death as compared to fast death, top differentially regulated cytokines, enzymes, peptidases, transcription regulators, and plasma membrane proteins were investigated. The top three upregulated cytokines in the skin of slow death individuals were *CXCL1, CCL23* and *SPP1* while the top three downregulated cytokines were *CXCL14, WNT4 and WNT2* (**Supplementary Fig. S2A-B**). The enzymes *GPX3*, *GALNT15* and *SULT1E1* were upregulated in skin cells of slow death-type, while *TYRP1*, *HSD3B1* and *TPPP* were downregulated (**Supplementary Fig. S2C-D**). The upregulated peptidases in skin of slow death-type included *MMP2, ADAMTS9* and *PCSK1*, while *RAB7B, MMP27* and *ENDOU* were downregulated (**Supplementary Fig. S2E-F**). At the level of transcription regulators, *ZBTB16, KLF25* and *CEBPD* were upregulated in skin cells of slow death-type, while *POU3F1, MYBL2* and *CYS1* were downregulated (**Supplementary Fig. S2G-H**). The plasma membrane proteins *CD79B*, *TLR4* and *TLR2* were upregulated in skin of slow death-type, while *SFRP2, FCER1A* and *TNFRSF18* were downregulated (**Supplementary Fig. S2I-J**). Investigation of pathways enriched in these transcripts in slow death using KEGG database identified PI3K-Akt signaling to be the top upregulated pathway in slow death skin samples (**Fig. 2A; Supplementary Table 4**). Of interest, specific pathways supporting functional properties of the skin as an organ were downregulated. Such pathways included melanogenesis, and those representing the skin extracellular matrix (collagen expression and metabolism) (**Fig. 2B; Supplementary Fig. S1E**). PI3K-Akt signaling is a developmental and survival pathway that is known to regulate cell differentiation, proliferation and apoptosis^16–18^. Twenty-five PI3K-Akt signaling pathway genes upregulated in skin cells from slow death are presented in **Fig. 2C-D**. Interestingly, these 25 genes were independent of post-mortem ischemic time and sex of the subjects (**Fig. 2E and Supplementary Fig. 3A-C)**. Additionally, FoxO signaling, and focal adhesion pathways were the other significant pathways upregulated in skin of slow death individuals (**Fig. 2A, 2D; Supplementary Table 4**). Interestingly, the 3 stemness genes *FOXO1*, *DLL1* and *PAF1,* which are annotated in stem cell population maintenance gene ontology term (EMBL-EBI; homo sapiens; GO:0019827), were increased in slow death skin compared to fast death indicating the appearance of stem cell features (**Supplementary Fig. S4A-C)**. To address the specific post-mortem cell compartment upregulating these developmental signaling, we turned towards single-nuclei RNA sequencing (snRNA-seq) dataset present in the GTEx database.

**Figure 2.**
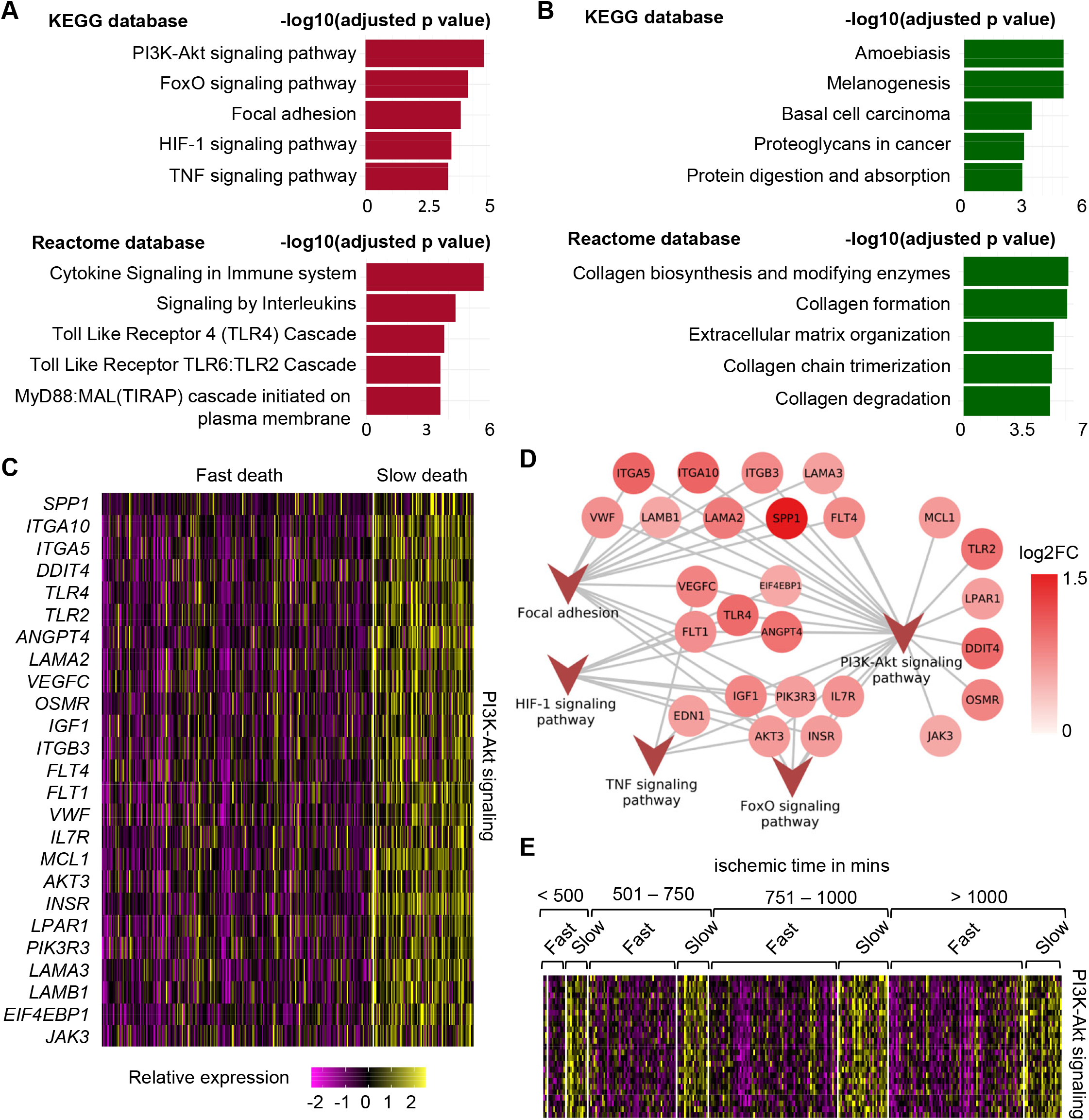
Upregulation of PI3K-Akt mTOR signaling in human post-mortem skin samples from subjects with longer terminal phase (slow death). (**A**) Top 5 upregulated pathways in skin cells of slow death-type cases compared to fast death cases using KEGG (upper) and Reactome (lower) databases. (**B**) Top 5 downregulated pathways in skin cells of slow death-type cases compared to fast death-type cases using KEGG (upper) and Reactome (lower) databases. (**C**) Heatmap showing upregulated genes involved in PI3K-Akt signaling pathway in slow death-type cases. (**D**) Network representing the identified genes in PI3K-Akt signaling pathways and their involvement (edges) in other upregulated pathways i.e., FoxO signaling pathway, focal adhesion, HIF-1 signaling pathway and TNF signaling pathway. (**E**) Heatmap representing PI3K-Akt signaling pathway genes (*panel C)* split by post-mortem ischemic time bins.

### Post-mortem skin fibroblast population

The single-nucleus RNA-seq (snRNA-seq) data from 5,327 post-mortem skin nuclei (n=3, GTEx database) were analyzed. Cluster analyses of these cells resulted in UMAP plot identifying six clusters with distinct expression profiles (**Fig. 3A).** These clusters comprised of five different cell types in human post-mortem skin: adipose (PLIN1^high^; 1.16%), endothelial (VWF^high^; 19.2%), epithelial (CDH1^high^; 52.96%), fibroblast (COL1A2^high^; 9.93%), immune (IL1R2^high^; 4.77%), and rest other uncharacterized compartments (11.98%) (**Fig. 3A-F**). PI3K-Akt signaling score was calculated in these five identified compartments which was upregulated in fibroblasts (log2FC = 0.53) followed by in endothelial cells (log2FC = 0.22) using the Wilcoxon Rank sum test (**Fig. 3G-H, Supplementary Table 5**). The genes *LAMA2*, *AKT3*, *LPAR1*, *INSR* and *LAMB1* have the highest expression average in fibroblasts (**Supplementary Table 5**). Akin to the bulk transcriptomic dataset (**Fig. 2E)**, PI3K-Akt signaling was also independent of post-mortem ischemia in snRNA-seq dataset **(Fig. 3I-J**).

**Figure 3.**
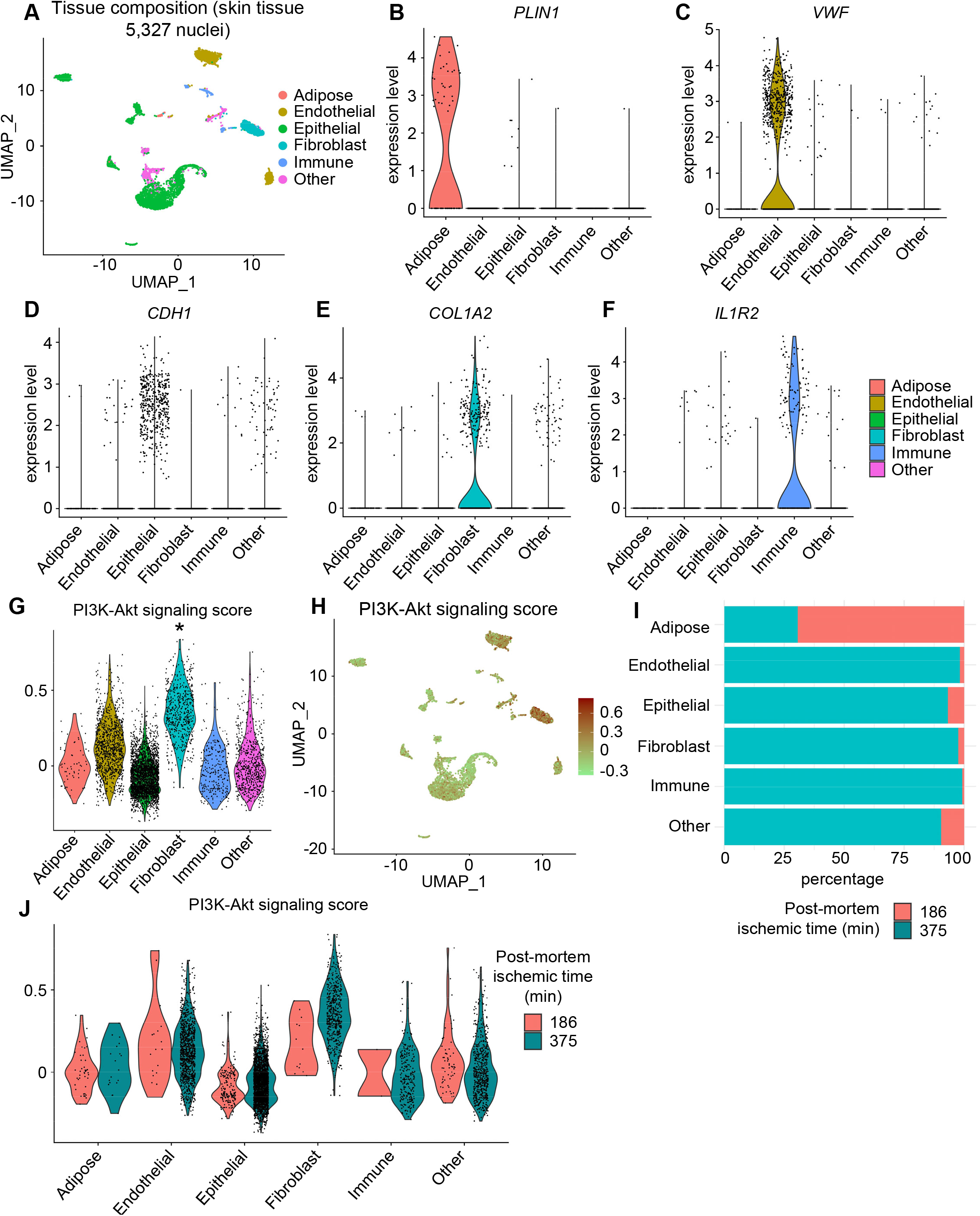
Single nuclei (sn)RNA-seq dataset identified the maximum enrichment of genes from PI3K-Akt signaling pathway in fibroblasts of human post-mortem skin. (**A**) UMAP plot showing cellular composition. (**B**) Violin plot showing the expression level of adipose (*PLIN1*^hi^), (**C**) endothelial (*VWF*^hi^), (**D**) epithelial (CDH1*^hi^*), (**E**) fibroblast (*COL1A2*^hi^), and (**F**) immune (*IL1R2^hi^)* compartments. Rest other minor cell types were grouped as *others*. (**G**) Violin plots and (**H**) UMAP plot representing PI3K-Akt signaling pathway score in different skin compartments mentioned in C-F. (*adjusted p value < 0.05 and fold change > 0.3). (**I**) Proportion of cells within each compartment with different PMI time points. Colors are same as in **J**. (**J**) PI3K-Akt signaling score in skin tissue compartments in different post-mortem ischemic (PMI) time points (in min).

### Transcriptional regulation of PI3K-Akt signaling in the post-mortem skin

To identify the upstream regulators of PI3K-Akt signaling genes found to be upregulated in skin cells from slow death-type samples, IPA software was used. The genes significantly enriched in PI3K-Akt signaling pathway (from bulk RNA-seq data) along with all upregulated transcription factors (log2FC>0.25), in slow death-type, were submitted in IPA to test direct functional relationships. The resulting network was visualized using CytoScape software^19^. This analysis of upstream regulators of PI3K-Akt signaling pathway identified *FOXO1*, *FOXO3*, *ATF4*, *CEBPB* and *CEBPD* to be among the master regulators for the pathway^20–23^ (**Fig. 4A**). Similar analysis using snRNA-seq dataset (**Fig. 3**), identified COL1A2^high^ fibroblast compartment to have the highest score for upstream regulators of PI3K-Akt signaling (**Fig. 4B-C**). Hence, fibroblasts from post-mortem skin not only upregulated genes of PI3K-Akt signaling, but they also displayed higher transcripts of the upstream regulators of this developmental pathway. This finding was supported at protein levels by a subsequent prospective study comprising of 16 post-mortem subjects (**Supplementary Table 6**). Compared to fast death subjects, the skin samples of slow death subjects showed remarkably elevated AKT protein levels in COL1A2+ fibroblasts (**Fig. 4D-E**). Interestingly, in addition to fibroblasts, post-mortem endothelial cells have higher scores for upstream regulators of PI3K-Akt signaling (**Fig. 4B-C**). Thus, active crosstalk between different cellular compartments of the post-mortem skin is expected. To understand such crosstalk, cell-cell interaction (CCI) was calculated among five different cellular clusters (e.g., endothelial, epithelial fibroblasts, immune and adipose cells) identified in snRNA-seq data using CellChat^24^. CellChat analysis resulted an aggregated connectome specifying the distribution of 62 cell-cell interactions (p<0.01, **Fig. 5A-B**). Fibroblasts emerged as the main compartment for initiating PI3K-Akt signaling as evident by the identification of laminin signaling ligands (*LAMA2*, *LAMB2*, *LAMC1*, *LAMC3*) (**Fig. 5A-D, Fig.2C**). CD44 was found to be the most predominant receptor for PI3K- Akt signaling and was found to be upregulated in fibroblasts, epithelial and immune cells (**Fig. 5C-D)**. These findings recognized fibroblasts in post-mortem skin to be a major hub for PI3K-Akt signaling pathway which is adequately supported by other cellular compartments by specific ligand-receptor signaling (**Fig. 5A-D, Fig.2C**).

**Figure 4.**
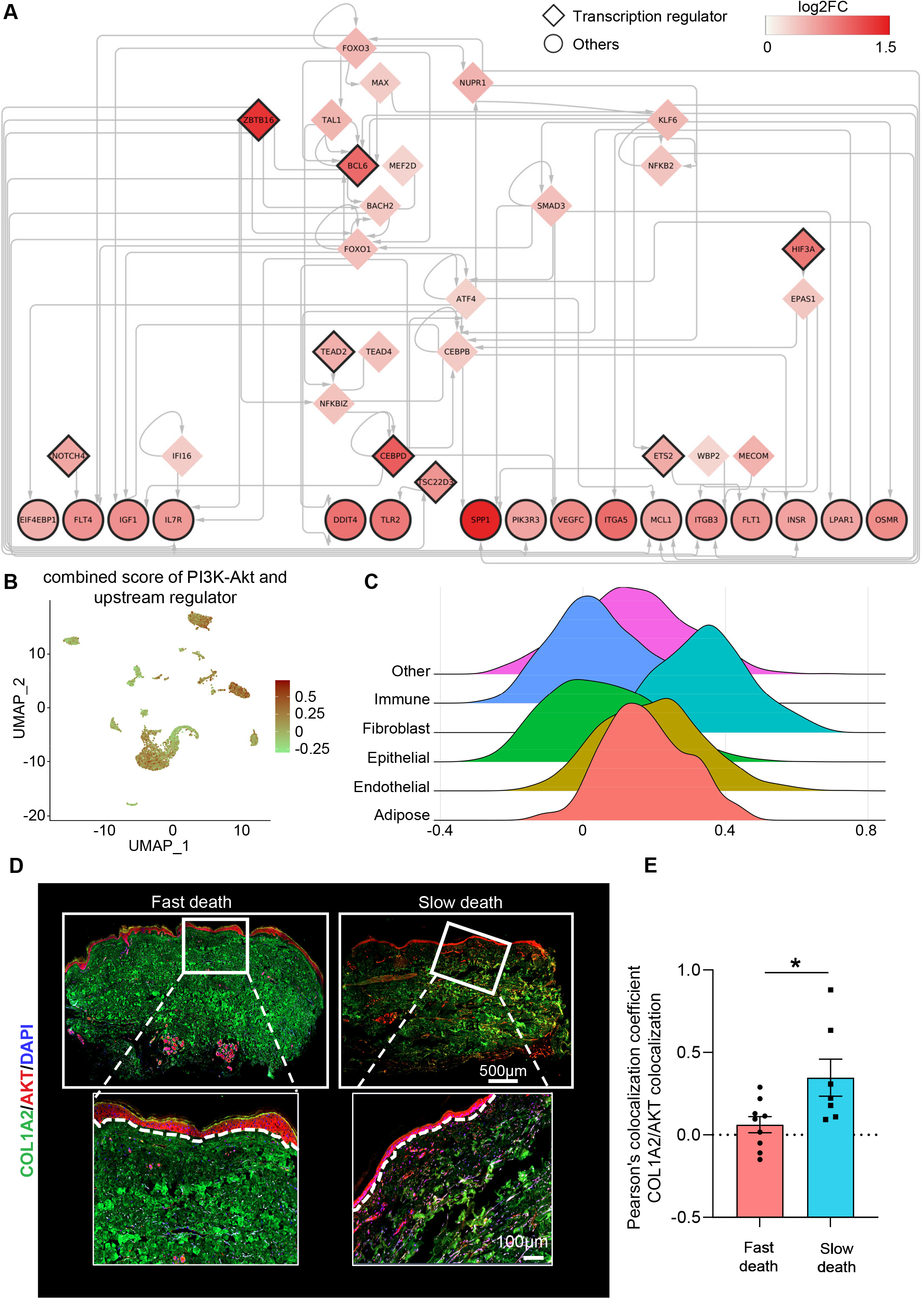
Upstream regulators of PI3K-Akt signaling pathway upregulated in human post-mortem skin samples from subjects with longer terminal phase (slow death). (**A**) Gene regulatory network representing upregulated genes involved in PI3K-Akt signaling pathways (ellipse-shaped nodes) along with their upstream transcription factors (diamond-shaped nodes) as computed using IPA. Nodes with black borders represent genes with log2FC > 0.5, while nodes without borders represent genes with log2FC between 0.25 and 0.5. **(B)** UMAP plot and **(C)** ridge plot of scores calculated for the genes upregulated in PI3K-Akt signaling pathway and their upstream regulators in snRNA-seq data. **(D)** Representative immunofluorescence analyses for COL1A2-AKT colocalization in lower limb skin from human post-mortem subjects. **(E)** Pearson colocalization coefficient calculation of COL1A2-AKT colocalization in skin tissue of slow death-type (Hardy scale 4) cases compared to fast (Hardy scale 1 and 2) death-type cases. (n = 9 and 7, *P < 0.05, Student’s t test). Data represented as mean ± SEM.

**Figure 5.**
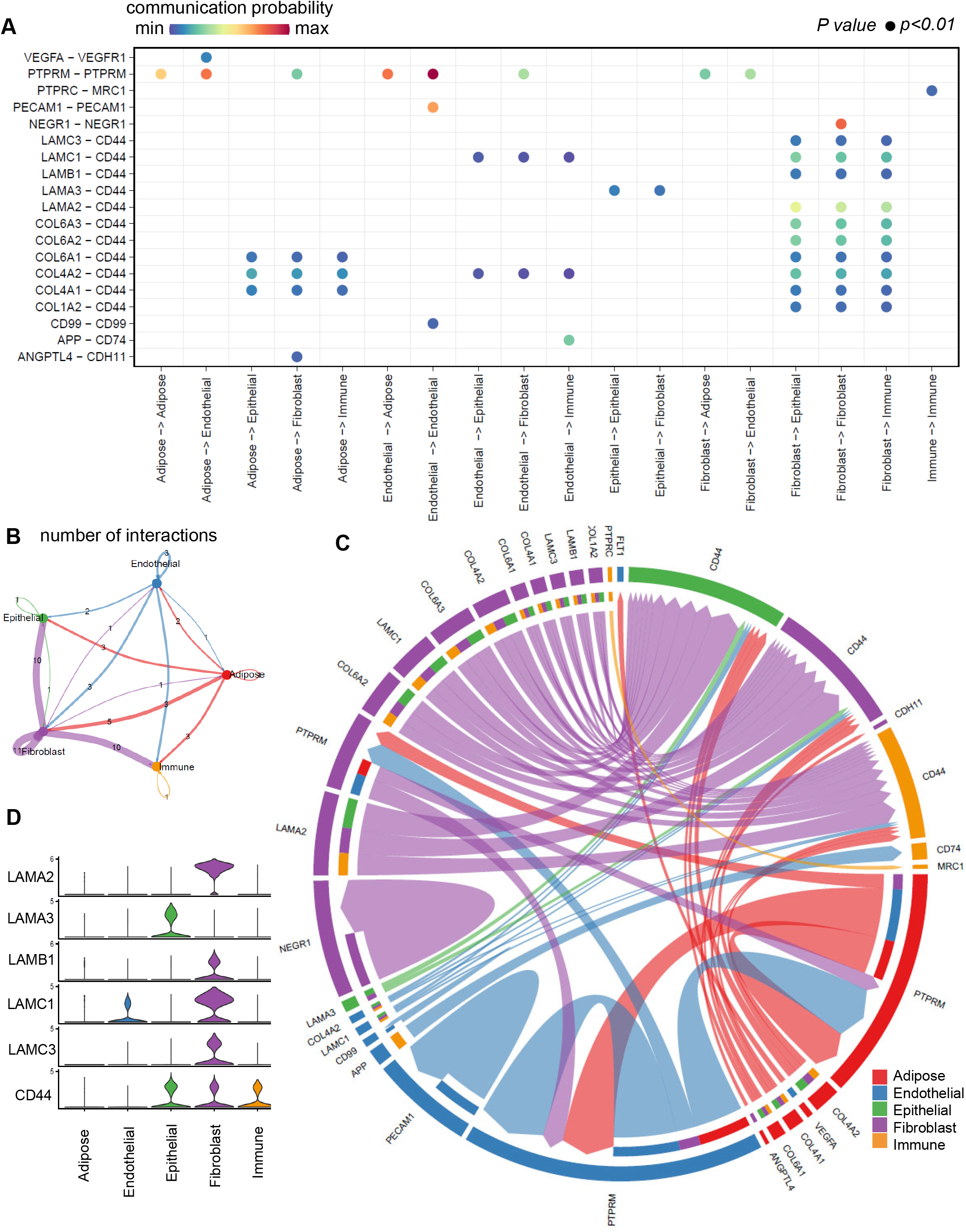
Cell-cell communication analyses between different compartments in human post-mortem skin identified fibroblast as the major signaling hub. (**A**) Bubble plot representing significantly enriched (p<0.01) cell-cell communication mediated by multiple ligand-receptors and/or signaling pathways. (**B**) CellChat analysis resulted an aggregated connectome specifying the distribution of 62 cell-cell interactions. (**C**) Chord plot representing ligand receptor pairs involved in each significant communication between pair of cell type. Relative strength of the signal is identified by the width of the chord. (**D**) Violin plot representing the ligand receptor pairs involved in PI3K-Akt signaling in the post-mortem skin. Fibroblasts were the major compartment involved in this signaling.

### Other significant biological processes in the post-mortem skin associated with the longer terminal phase

Further characterization of upregulated biological processes in skin cells of slow death-type samples included vasculature development, blood vessel formation, angiogenesis, and endothelial cell migration (**Fig. 6A**). On the other hand, extracellular matrix organization, collagen metabolic process and antigen processing and presentation were downregulated in the skin of slow death-type samples (**Fig. 6B**). The upregulated genes in slow death-type involved in angiogenesis process included *PROK1*, *ANGPTL4* and *GLUL* among others (**Fig. 6C)**. Similar to the development pathways (**Fig. 2E)**, upregulation of angiogenesis in skin cells of slow death-type cases were neither found to be dependent on ischemic time nor on the sex of the individual (**Supplementary Fig. 4B-C**). The upregulated growth factors in slow death with established role in angiogenesis were *ANGPT4*, *EPGN*, *PROK1* and *VEGFC* (**Supplementary Fig. S5A**). Other growth factors identified included *GDF15*, *PROK1*, *EDN2*, *DKK1* to be upregulated in skin of slow death-type cases (**Supplementary Fig. S5A**). The bulk RNA seq findings were then complimented with the snRNA-seq analysis to identify the cellular compartments involved. As expected, snRNA-seq of skin cells identified the endothelial compartment to have the highest level of angiogenesis score (log2FC = 0.42) (**Fig. 6D-G**). The genes *PTPRM*, *AKT3*, *CLDN5*, *HIF3A* and *FLT1* have the highest expression average in endothelial cells among the genes that were used to calculate the angiogenesis score (**Supplementary Table 5**). Post-mortem endothelial cells were the major receiver (*FLT1*), while adipose cells (*VEGFA*) primarily contributed to the outgoing VEGF signaling (**Fig. 5C, Supplementary Fig. S5B-C)**. Interestingly, skin fibroblasts also contributed to angiogenesis (log2FC = 0.22 compared to other skin compartments) (**Fig. 6D-G**). This observation is consistent with the recently reported notion of the vasculogenic fibroblasts^25^. PTPRM signaling was predominantly responsible for such vasculogenic behavior of the post-mortem fibroblasts. Taken together, these findings recognize the presence of specific cellular connectome responsible for the observed increase in developmental genes in the post-mortem skin of slow-death subjects with longer terminal phase.

**Figure 6.**
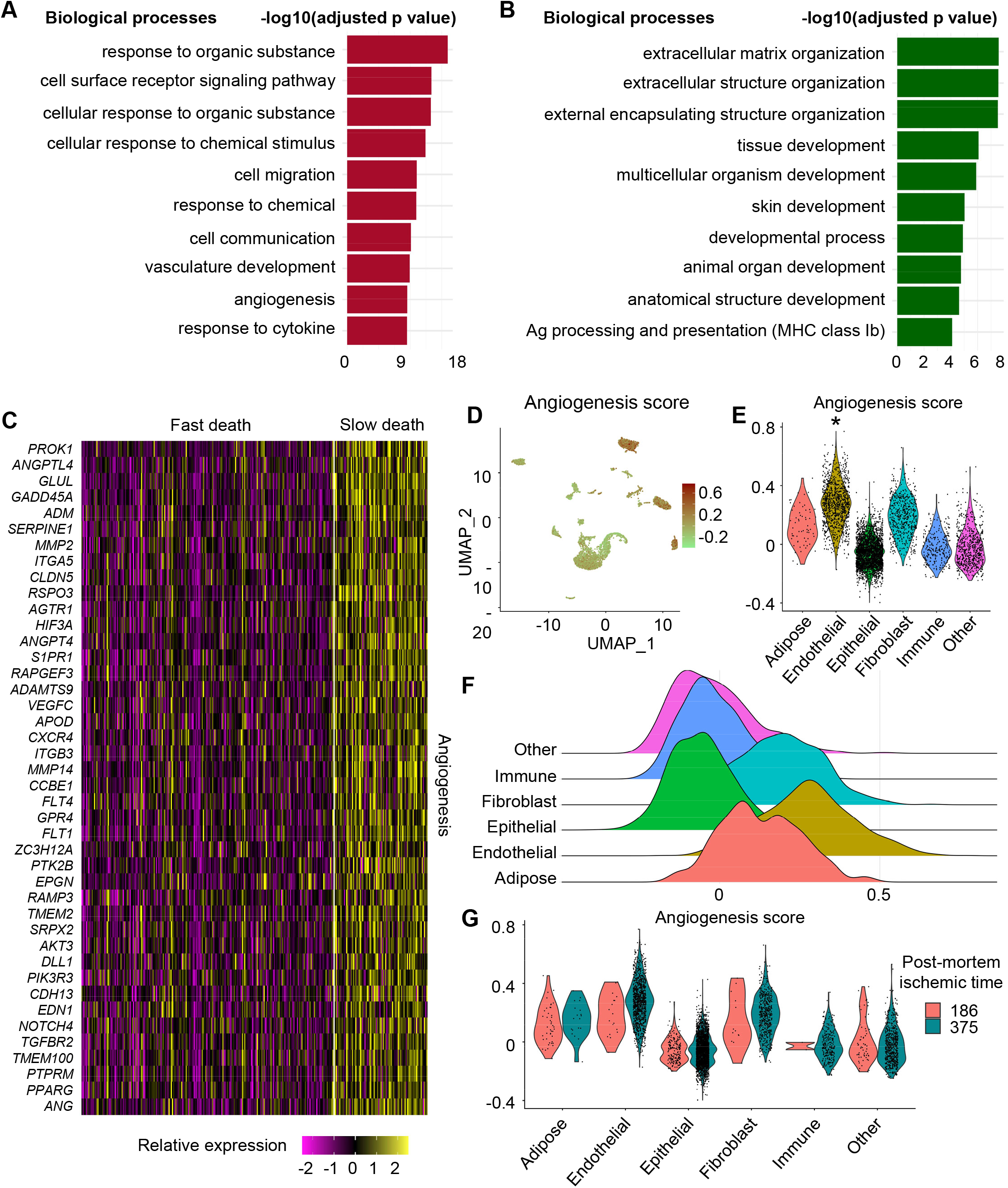
Upregulation of angiogenesis biological process in human post-mortem skin samples from subjects with longer terminal phase (slow death). (**A**) Top 10 upregulated and (**B**) downregulated biological processes in post-mortem skin of slow death-type cases compared to fast death-type cases. (**C**) Heatmap representing relative expression of upregulated genes involved in angiogenesis in slow death-type cases. (**D**) UMAP and (**E**) Violin plots representing angiogenesis score in different skin tissue compartments. (*adjusted p value < 0.05 and fold change > 0.3). **(F**) Ridge plot representing angiogenesis score distribution in different skin tissue compartments. (**G**) Angiogenesis score in skin tissue compartments in different post-mortem ischemic time points (in min).

## Discussion

Human organismal death initiates a state within the body wherein certain tissue components strive to survive by mounting the expression of developmental genes. This phenomenon offers an opportunity to gain insight into death as a biological variable. Of outstanding interest is our observation that in humans such survival response is mounted following “slow” but not “fast” (traumatic) death. The terminal phase of life initiates a state within the body in which individual cells within tissues strive to survive by adjusting their transcriptional program thus upregulating developmental pathways such as PI3K-Akt signaling. Such slow-death post-mortem signaling is further supplemented with upregulation of embryonic development related genes such as ATF4 and FOXO. The PI3K-Akt survival pathway also regulates distinct physiological activities such as cell differentiation and proliferation^16–18^. Furthermore, Akt inhibits apoptosis and promotes skin regeneration^26, 27^. Upregulation of FoxO signaling in slow-death post-mortem skin is likely to contribute to cell survival. Different FoxO family genes are known for their regenerative potential in stem cells^28^. Unsurprisingly, although limited, sex-related difference were observed in response to slow death-type compared to fast death-type samples. Specifically, *YTHDC1* was upregulated in females in slow death-type cases and not in males. *YTHDC1* increases Akt phosphorylation in ischemic tissue and facilitate neuronal survival^29^. Another gene, Serum Amyloid A4 (SAA), found to be upregulated in female slow death-type cases, also activates PI3K signaling^30^ and angiogenesis pathways^31^. The selective mounting of multiple survival pathways in the post-mortem skin unveils a novel paradigm recognizing death itself as a trigger of mechanisms that must be accounted for while interpreting experimental data from tissues harvested from the dead.

Contrasting results in slow versus fast death scenarios underscore the critical importance of the type of death in question with emphasis on the terminal phase of human life. Upstream analysis of PI3K-Akt signaling pathway identified key regulators such as *CEBPD*, *ATF4*, *CEBPB*, *FOXO3* and *FOXO1*. *ATF4* has been previously reported to possess survival and anti-apoptotic roles^20, 21^. Besides their role in survival, *FOXO1* and *FOXO3* regulate embryonic development^32, 33^. Further investigation of distinct tissue compartments using snRNA-seq data revealed that PI3K-Akt signaling has highest activity in skin fibroblast compartment. In the PI3K-Akt signaling pathway, upstream regulators such as *LAMA2*, *AKT3*, *LPAR1*, *INSR* and *LAMB1* were highly expressed in post-mortem dermal fibroblasts^34–36^. Developmentally, AKT1 is implicated in the differentiation of multiple cell types^37^. Overexpression of AKT1 was reported to increase self-renewal and differentiation of stem cells in cortical progenitors^38^. In the skin, blocking of PI3K-Akt pathway resulted in suppression of self-renewal of skin-derived precursors. Activation of this pathway alleviated such senescence response and promoted cell proliferation^39^. Additionally, PI3K-Akt signaling protected melanocytes from oxidative damage. Downregulation of this signaling caused melanocyte apoptosis^40, 41^. Of note, downregulation of AKT inhibit Epithelial-to-Mesenchymal (EMT) process^42, 43^. The EMT process is necessary for skin wound repair^44–46^.

Angiogenesis, blood vessel development and tube morphogenesis were other significant pathways enriched in skin cells of slow death-type cases. snRNA-seq data revealed that endothelial skin compartment has the highest score for angiogenesis in post-mortem skin tissue. *PTPRM*, *AKT3*, *CLDN5*, *HIF3A* and *FLT1* have the highest expression values in post-mortem endothelial cells among the genes that were used to calculate the angiogenesis score. Induction of angiogenic gene expression is known to be triggered by hypoxia^47^. Slow death is known to be associated with prolonged systemic mild-to-moderate hypoxia caused by immobility and other factors^48^. During this time, tissues remain viable and at a temperature conducive to gene expression thus explaining upregulation of the hypoxia-inducible PI3K-Akt pathway^49^. In fast death, frequently associated with sudden and rapid loss of blood supply and core body temperature, bodily tissues approach anoxia and hypothermia not conducive for efficient gene expression. Indeed, several reports have linked HIF1α and its stabilization as a result of hypoxic pressure to regenerative processes in mammalian tissues^50^, and in humans these processes have been used to facilitate wound healing^51^ and healthy longevity^52^. For such life-supporting adaptive processes to be induced in a post- mortem setting could of functional significance in settings of organ transplantation.

Efforts to understand the potential of death as a biological variable (DABV) in influencing the transcriptomic composition of surviving component tissues has far-reaching significance including interpretation of experimental data collected from the dead and mechanisms involved in transplanted organs obtained from dead donors. Activation of survival and angiogenic pathways in sun-exposed skin samples from slow death-type compared to fast death-type revealed the effect of the prolonged terminal phase of life on post-mortem transcriptional program changes. Gene expression alteration during the terminal phase of life, and its functional significance, needs further investigation to identify how cells activate distinct transcriptional programs in an attempt for survival which might reveal novel directions in regenerative medicine discipline. Discovering how to turn on and off death-induced tissue reprogramming in the live could give people access to life saving interventions, correct genetic errors, and restore functionality that was thought lost, ushering in a transformative change to the current paradigm of experimental biology.

## Methods

### Data download

Gene expression (transcripts per million) TPM values and sample annotations, including the duration of terminal phase of life, were downloaded from GTEx database (accession phs000424.v8.p2)^11^. Out of the 17,382 samples available in GTEx portal, 701 sun-exposed human skin samples were extracted and analyzed. Gene expression values were log normalized and PCA (Principal Component Analysis) was performed for dimensionality reduction. Next, Uniform Manifold Approximation and Projection (UMAP) was applied for sample visualization using Seurat^53–57^. Scater package in R was used to identify the percentage of variance explained by ischemic time, RNA quality, age, sex and death-type among sun-exposed skin samples^58^.

### Data analysis

The 701 samples were classified based on the 4-point hardy scale as provided in GTEx^11^. Differential gene expression analysis was performed using the Wilcoxon rank-sum test to compare between sun-exposed skin samples resulting from slow death type (>24h terminal phase; 81 samples) and fast death type: violent or unexpected death (<1h terminal phase; 220 samples). The protein coding genes based on UniProtKB/Swiss-Prot database were kept for downstream analysis^59^. STRING database^60, 61^ was used to retrieve both functional and physical protein associations using the differentially expressed genes (DEGs) with |log2FC| > 0.5 with a medium confidence score (0.4) as reported by us previously^46, 62^. Furthermore, genes with adjusted p value < 0.01 and log2FC ± 0.5 were chosen to perform the function enrichment analysis using g:Profiler with default parameters^63^. DEGs were investigated in Ingenuity Pathway Analysis (IPA) software as reported by us^64–66^ to further classify them based on their functional class including transcription factor, growth factor, cytokines and transmembrane receptors^67^. To identify whether there is a sex-based response in slow death-related gene expression change or not, slow and fast death cases were compared in males and females separately. In male, 47 slow death and 168 fast death samples were compared, and 34 slow death and 52 fast death samples were compared in females. Wilcoxon Rank Sum test was used to identify DEGs between slow and fast death samples.

### Pathway analysis

To identify which skin compartment has higher score for PI3K-Akt signaling and angiogenesis, snRNA-seq data from GTEx database was retrieved. Skin samples were selected, including 5,327 nuclei, for downstream analysis using Seurat package in R^68^. Gene expression values were log normalized with 10,000 as a scaling factor. Then, the top 2,000 variable genes between skin nuclei were identified and scaled. Next, principal component analysis was performed, and the first 20 principal components were chosen for visualization using UMAP. Tissue compartments annotation data was retrieved from the sample metadata file provided by GTEx database. Then, PI3K-Akt signaling, and angiogenesis scores were calculated for each nuclei using the identified upregulated genes from the bulk data analysis using the function *AddModuleScore* in Seurat. The function compares the gene expression average of each gene set with a random control gene set to calculate a score for the desired set of genes.

To identify significant incoming and outgoing cell-cell interaction among different skin compartments, CellChat package in R was used to identify over expressed ligands and receptor across cell skin compartments^24^. The database in CellChat included 1,199 Secreted Signaling, 421 ECM-Receptor and 319 Cell-Cell Contact interactions.

### Study Approval

Since the skin samples investigated in this study were collected from deceased individuals, IRB of Indiana University decided that this does not meet the regulatory definition of ‘human subject research,’ i.e., “a living individual…”.

### Histology and Immunohistochemistry

Histology of post-mortem skin was performed from 10- μm-thick paraffin sections as reported previously^69–72^. Paraffin tissue sections were deparaffinized, blocked with 10% NGS and incubated with primary antibody against AKT (cat#9272, 1:200 dilution) and COL1A2 (cat#sc-393573. Santa Cruz. 1:200) followed by appropriate fluorescence conjugated secondary antibodies (Alexa 568-tagged for AKT, 1:200 dilution; Alexa 488-tagged for COL1A2, 1:200 dilution). Images were collected using a Zeiss Axio Scan Z1 guided by Zen blue imaging software and colocalization was calculated using Zen blue software^73–75^.

## Acknowledgment

This work was supported by John Templeton foundation grant ID-61742. This work was also supported in part by National Institute of Diabetes and Digestive and Kidney Diseases (NIDDK) grants DK128845 and DK125835 to CKS. This work was also supported in part by U.S. Department of Defense grants W81XWH-21-1-0033 to CKS and W81XWH-22-1-0146 to KS. Research programs led by CKS was supported by the Lilly Endowment INCITE (Indiana Collaborative Initiative for Talent Enrichment) program.

## Authors’ contribution

AA, KS and CKS designed the study. AA collected the dataset from public domain and imple- mented the available tools for sequencing data analysis with KS, RS, JW and SL. KS, SS, XG, SK, GS, AH, SR, and CKS participated in post-mortem skin collection and related data analysis. AA, and KS wrote the manuscript. JW and CKS thoroughly reviewed and edited the manuscript. All authors read and approved the final manuscript.

---

**Supplementary Figure 1.**
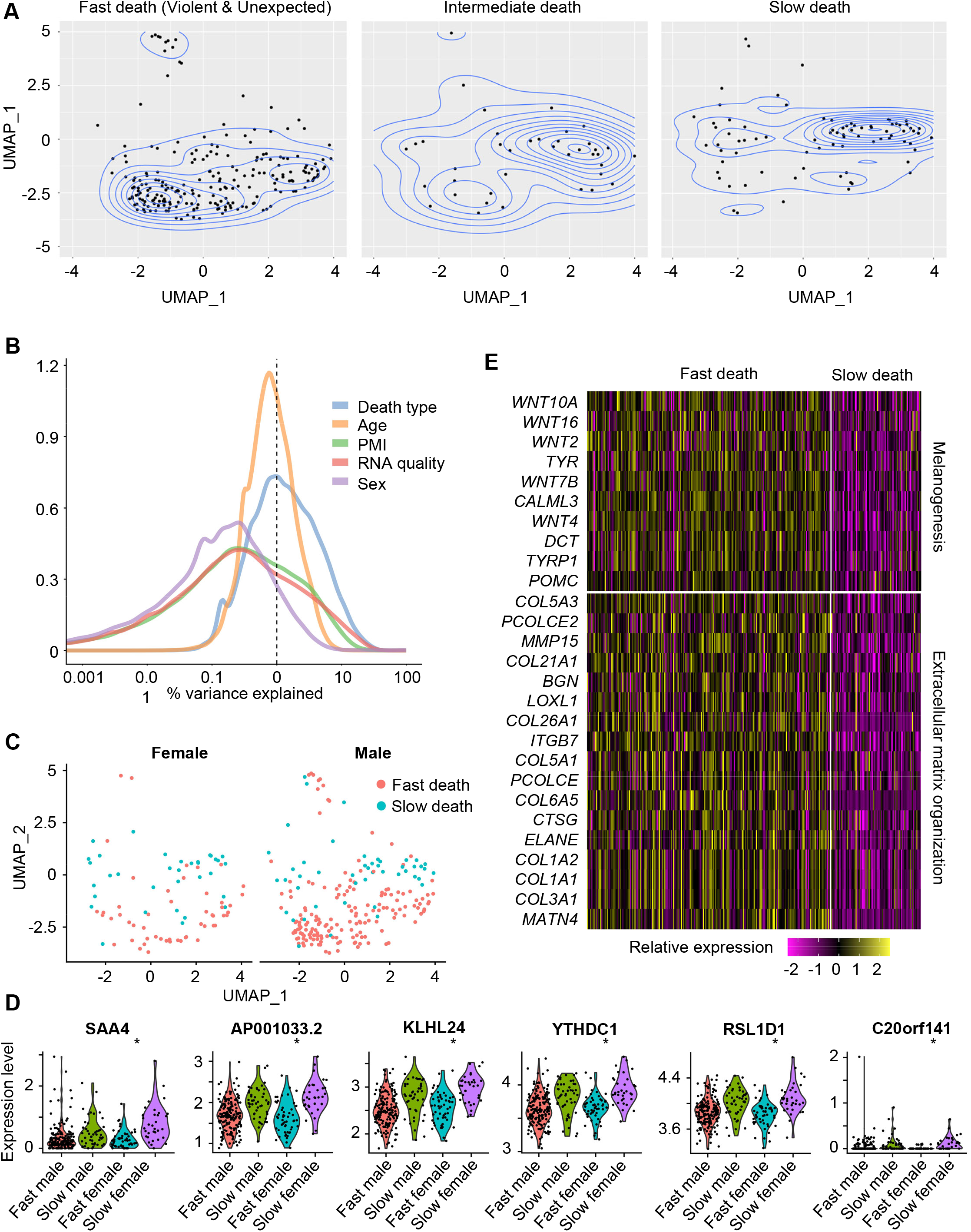
Variance explained by death type on Hardy scale. (**A**) Density plots for fast violent and unexpected death-type samples (left), intermediate death-type samples (middle) and slow death-type samples (right). (**B**) Variance explained by death type, age, post-mortem ischemia (PMI) time, RNA quality and sex among sun-exposed skin samples. (**C**) UMAP plots representing slow and fast death type samples in females and males. (**D**) Violin plots of the top genes having statistical significance (adjusted p < 0.05) to be upregulated in slow death in females and not in males. (**E**) Heatmap representing downregulated genes in skin of slow death-type cases compared to fast death-type cases involved in melanogenesis and extracellular matrix organization.

**Supplementary Figure 2.**
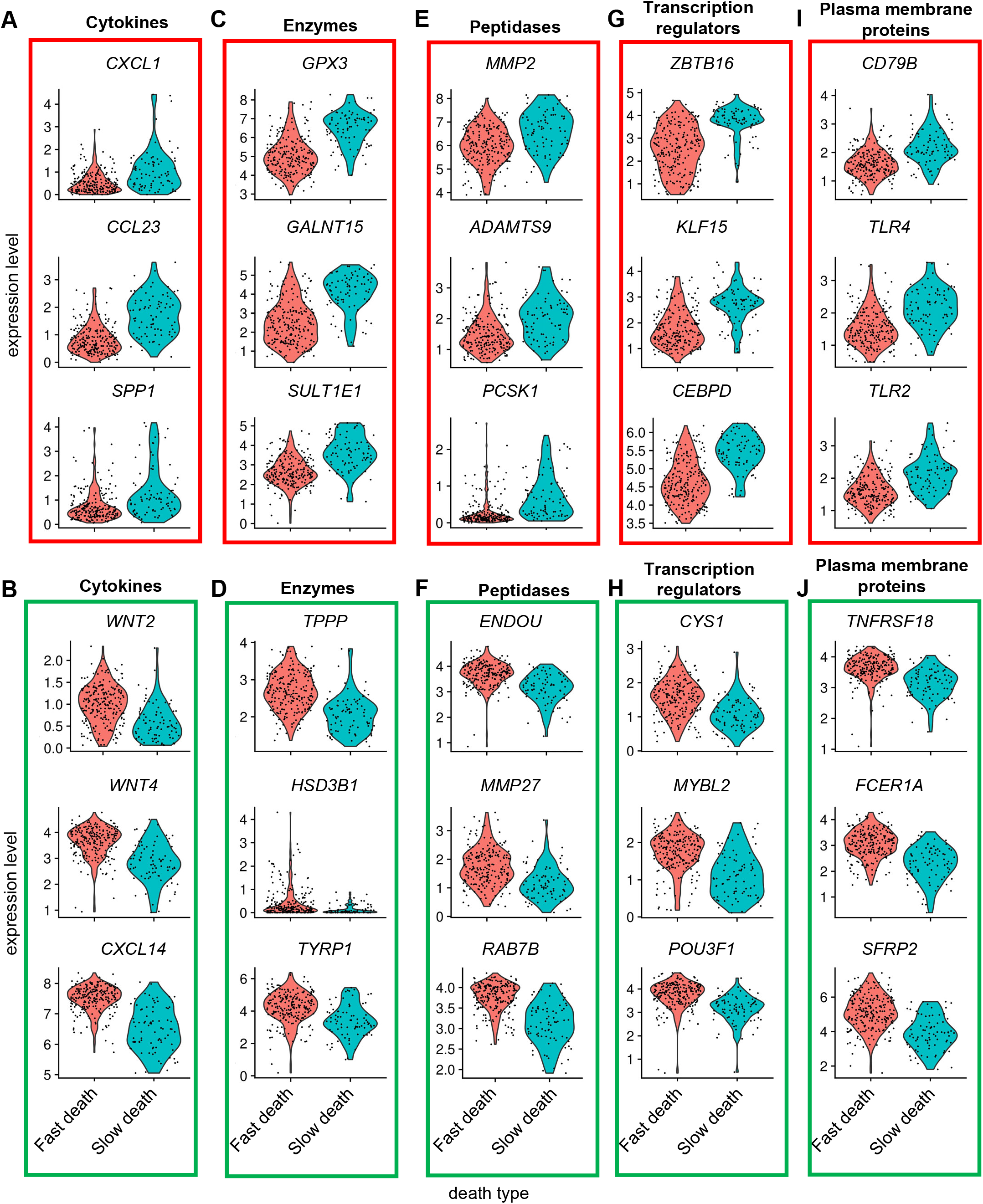
Expression level of top 3 upregulated and downregulated genes (adjusted p value <0.05) in post-mortem skin samples of subjects with slow death type. (**A-J**) Violin plots showing expression level of the top 3 upregulated and downregulated cytokines (**A-B**), enzymes (**C-D**), peptidases (**E-F**), transcription regulators (**G-H**) and plasma membrane proteins (**I-J**).

**Supplementary Figure 3.**
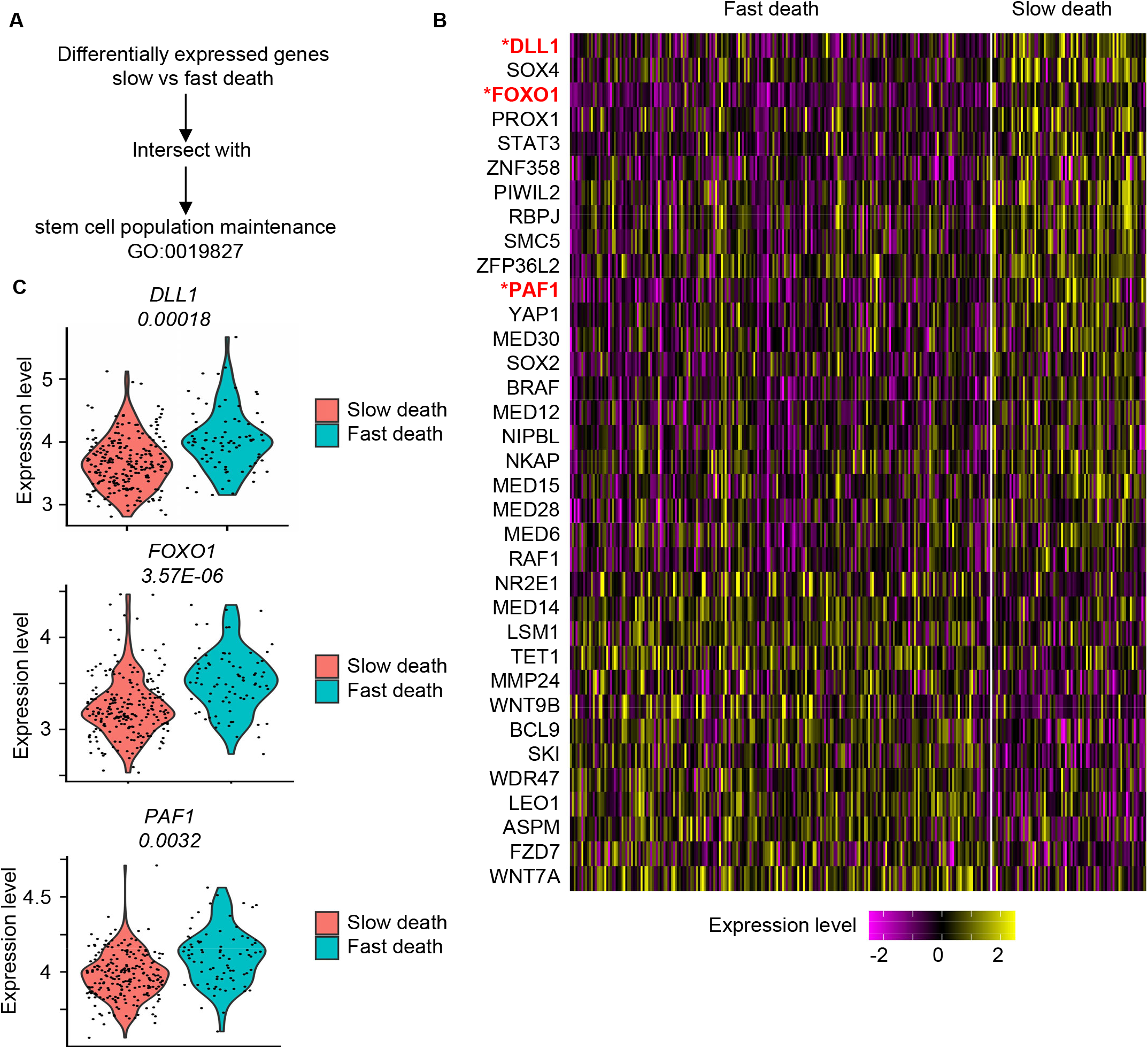
Upregulation of specific gene members of stem cell population maintenance. **(A)** The intersection between the differentially expressed genes when comparing slow death with fast death skin samples and the stem cell population maintenance biological process in homo sapiens (GO:0019827). (B) Heatmap representing differentially expressed genes having role in stem cell population maintenance with p value < 0.01 (genes with adjusted p value < 0.01 were marked with stars). **(C)** Expression level of the 3 identified genes having role in stem cell population maintenance with adjusted p value < 0.01.

**Supplementary Figure 4.**
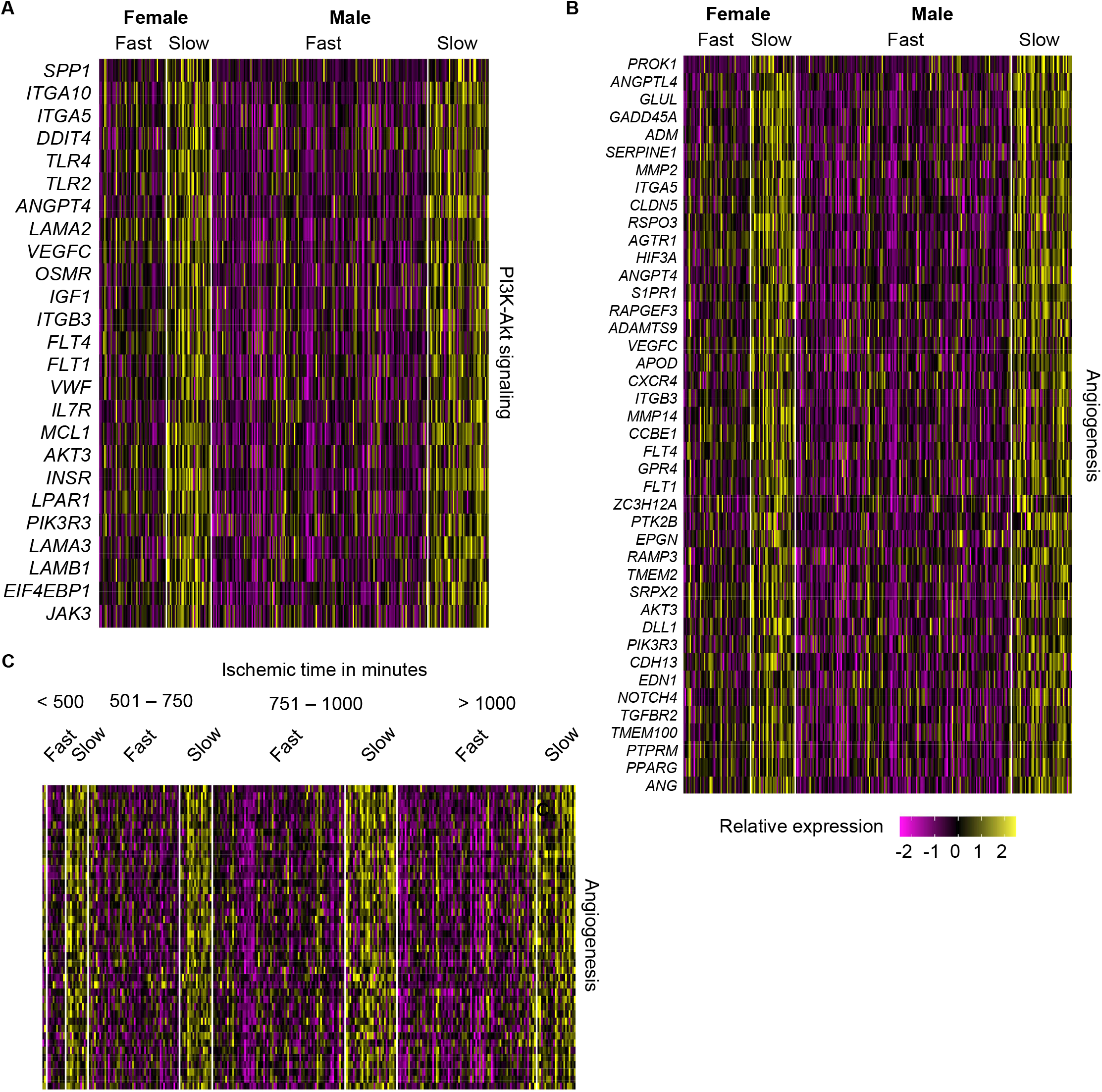
Sex-related response in human post-mortem skin in slow death on Hardy scale. (**A**) Heatmap representing relative expression of upregulated genes involved in PI3K-Akt signaling pathway as in Fig. 2C in male and female cases separately. (**B**) Heatmap representing relative expression of upregulated genes involved in angiogenesis as in Fig. 6C in male and female cases separately. (**C**) Same genes as in B (angiogenesis) separated by post-mortem ischemia time bins.

**Supplementary Figure 5.**
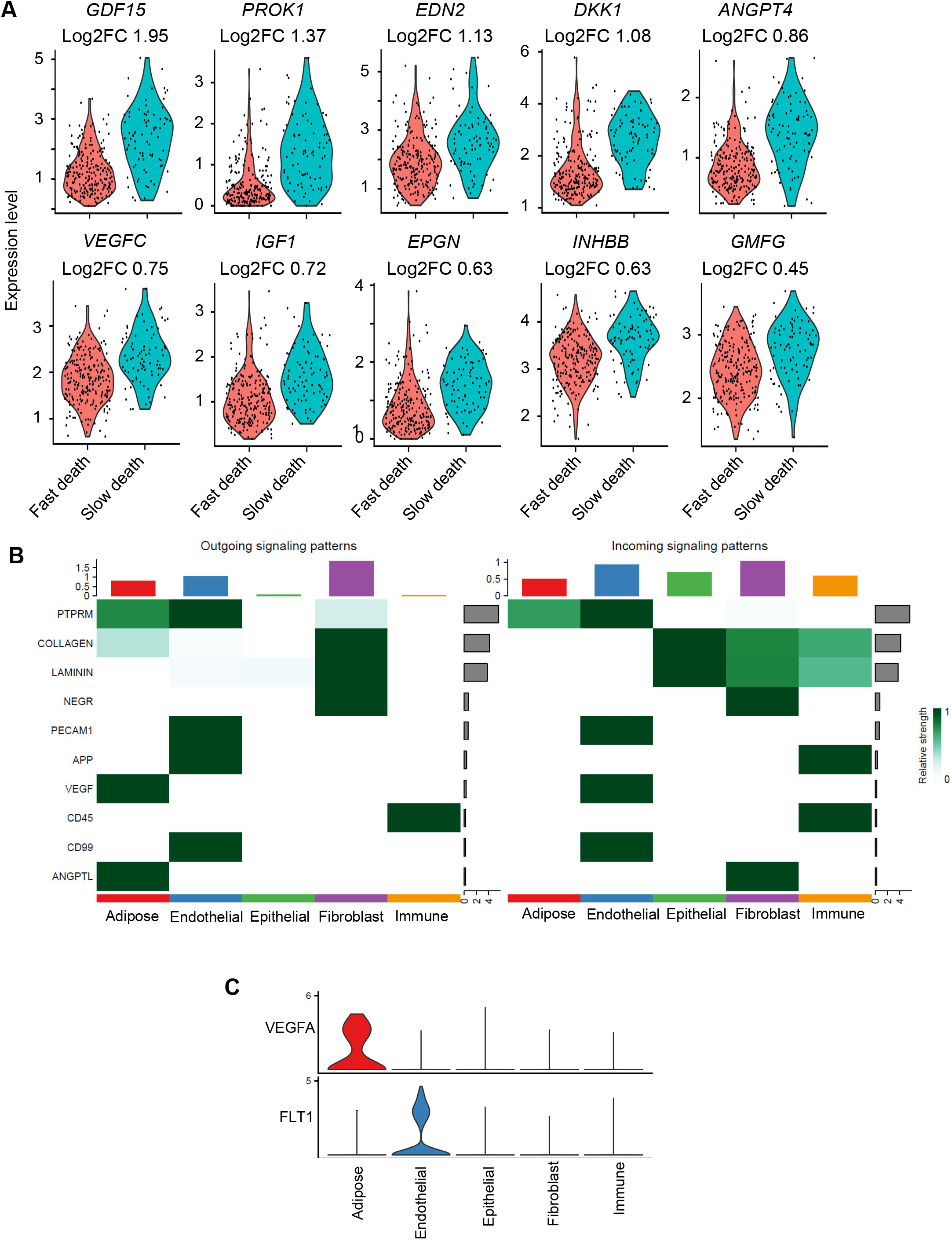
Expression level of differentially expressed growth factors in post-mortem skin samples based on death type. **(A)** Violin plots showing expression level of growth factors in skin samples of slow death-type cases compared to fast death-type cases (log2FC>0). Adjusted p value for the comparison between slow and fast groups < 0.01. (**B, C**) Heatmap and violin plots showing relative strength of the identified signals along incoming and outgoing signaling pattern among the cell types. Adipose cells were the major hub for outgoing angiogenic signal (*VEGFA*) while endothelial cells were predominant receiver (*FLT1*) of the angiogenic signals in the post-mortem skin.

**Supplementary Table 1. (Provided as excel sheet):** Sample annotations of 701 sun-exposed human skin samples that were downloaded from GTEx database (accession phs000424.v8.p2).

**Supplementary Table 2. (Provided as excel sheet):** Table showing 1,582 differentially expressed genes (DEG, adjusted p value < 0.01) as obtained from comparing gene expression level from slow death-type cells (Hardy scale score = 4) versus fast death-type samples (a hardy scale score of 1 and 2) (See sheet Significant DEG). Among them, 344 genes were upregulated, and 183 genes downregulated (cutoff log2FC >± 0.5). Among the identified DEG, a significantly enriched protein-protein interaction (PPI) network of 451 genes (nodes) was identified (see sheet Network analysis) connected with 1,892 edges (p value <1.0e-16) (see sheet STRING network).

**Supplementary Table 3. (Provided as excel sheet):** Comparison between males and females with respect to death type identified a total of 100 DEG (48 upregulated and 52 downregulated, see sheet common in M & F) in slow death samples irrespective of the sex (adjusted p value <0.05 and log2FC cutoff >±0.5).

**Supplementary Table 4. (Provided as excel sheet):** Investigation of pathways enriched in these transcripts in slow death using KEGG database revealed PI3K-Akt signaling to be the top upregulated pathway in slow death skin samples (see sheet KEGG). Additionally, FoxO signaling, and focal adhesion pathways were other significant pathways upregulated in skin cells from slow death individuals (see sheet KEGG).

**Supplementary Table 5. (Provided as excel sheet):** Table showing scores for PI3K-Akt signaling in five identified compartments in snRNA-seq dataset. Maximum enrichment of genes participating in PI3K-Akt signaling found to be upregulated in fibroblasts (log2FC = 0.53) followed by endothelial cells (log2FC = 0.22) (see sheet PI3K and angio scores). The genes LAMA2, AKT3, LPAR1, INSR and LAMB1 contributed the most in calculating PI3K- Akt signaling score in fibroblasts (see sheet PI3K avg). Next, genes PTPRM, AKT3, CLDN5, HIF3A and FLT1 contributed the most in calculating angiogenesis score in endothelial cells (see sheet angiogenesis avg).

**Supplementary Table 6.** Demographics of subjects for prospective validation studies

|  | Fast death (Hardy scale 1 and 2) | Slow death (Hardy Scale 4) |
| --- | --- | --- |
| n | 9 | 7 |
| Age (± SEM) | 57.2 ± 4.4 years | 71.9 ± 4.3 years |
| Sex | Male = 6 ; Female = 3 | Male = 3; Female = 4 |
| Post-mortem ischemia (± SEM) | 24.0 ± 3.9 hours | 19.5 ± 3.9 hours |
| autolysis score | none = 8; mild = 1 | none = 5; mild = 2 |
| Ethnicity | Caucasian = 3; African American = 6 | Caucasian = 3; African American = 4 |

